# Single-cell activity in human STG during perception of phonemes is organized according to manner of articulation

**DOI:** 10.1101/552315

**Authors:** Yair Lakretz, Ori Ossmy, Naama Friedmann, Roy Mukamel, Itzhak Fried

**Author notes:** Co-first authorship. Co-senior authorship (Roy Mukamel, Itzhak Fried). **Contact Information** Yair Lakretz, Cognitive Neuroimaging Unit, NeuroSpin Center, Gif/Yvette, France.

## Abstract

A long-standing controversy persists in psycholinguistic research regarding the way phonemes are coded in human auditory cortex during speech perception. Whereas the motor theory of speech perception suggests that phonemes are organized in terms of common articulatory gestures that generate them, auditory theories argue that phonetic processing is organized based on common spectro-temporal patterns in phoneme waveforms. Here, we recorded spiking activity in the superior temporal gyrus (STG) from six neurosurgical patients who performed a listening task with phoneme stimuli. Using a Naïve-Bayes model, we show that single-cell responses to phonemes are governed by articulatory features that have acoustic correlates (manner-of-articulation) and organized according to sonority, with two main clusters for sonorants and obstruents. We further find that ‘neural similarity’ (i.e. the similarity of evoked spiking activity between pairs of phonemes), is comparable to the ‘perceptual similarity’ (i.e. how much the pair of phonemes sound similar) based on perceptual confusion assessed behaviorally in healthy subjects. Thus phonemes that were perceptually similar, also had similar neural responses. Our findings establish that phonemes are encoded according to manner-of-articulation, supporting the auditory theories of perception, and that the perceptual representation of phonemes can be reflected by the activity of single neurons in STG.

## 1 Introduction

How are phonemes encoded in human auditory cortex during speech perception? Auditory theories argue that phonetic processing depends directly on properties of the auditory system (Jakobson et al. 1951; Stevens 1972, 1989, 2002). That is, listeners identify patterns in phoneme waveforms and match them with stored abstract acoustic representations. For example, vowels are characterized by a roughly bimodal spectra and sibilant fricatives by high-frequency energy. This view is consistent with early observations from language acquisition. During language development, manner-of-articulation distinctions are acquired early during childhood, compared to the place-of-articulation ones (Jakobson 1968; Grodzinsky and Nelken 2014). Since manner-of-articulation but not place-of-articulation features have identifiable acoustic correlates, this finding is consistent with auditory theories.

Alternatively, the motor theory of speech perception (Liberman et al. 1967; Liberman and Mattingly 1985) describes phoneme perception in terms of the articulatory gestures that generate it. According to this theory, the objects of speech perception are the intended phonetic gestures of the speaker, such as, ‘lip rounding’, or ‘jaw raising’. For example, the phoneme [m] consists of a labial stop gesture combined with a velum lowering gesture. The motor theory arose from an early observation that phoneme percepts are invariant across different contexts (Cooper et al. 1952; Liberman *et al.* 1967). In the case of co-articulation, several gestures overlap in time, which may cause the acoustic waveform of the same intended gesture to be significantly different than when it is pronounced in isolation. Therefore, a particular gesture can be represented by different acoustic waveforms in different phonemic contexts. Additional variation exists in the acoustic signal due to inter-speaker variability. This considerable variability led the supporters of the motor theory to propose that the objects of speech perception are not to be found in acoustics.

Phoneme perception has been the focus of numerous neuroimaging studies (Binder et al. 2000; Dehaene-Lambertz et al. 2005; Liebenthal et al. 2005; Möttönen et al. 2006; Desai et al. 2008; Formisano et al. 2008; Liebenthal et al. 2010; DeWitt and Rauschecker 2012) showing activation in regions that are selective to speech, over non-phonemic contrasts. Findings describe a hierarchical organization of regions in the temporal lobe from primary auditory and early posterior auditory areas processing low-level auditory features, to the anterior, ventral Superior Temporal Gyrus (STG) and Superior Temporal Sulcus (STS), processing higher-level phonemic features.

Electrocorticogram (ECoG) research of speech perception shows that the organization of phonemes can significantly differ across brain regions and tasks, depending on whether speech is being produced or perceived (Bouchard et al. 2013; Cheung et al. 2016). Bouchard et al. (Bouchard *et al.* 2013) showed that during production, phonemes in the ventral sensory-motor cortex (vSMC) are predominantly organized by place-of-articulation features (e.g., labial, alveolar, velar and glottal), while during listening, the organization was found to be dominated by manner features (Cheung *et al.* 2016). The same studies also showed that the dominant organizing feature in the STG during perception is also manner-of-articulation. Additionally, Pasley and colleagues showed that waveforms reconstruction from local field potentials in the lateral STG is highest for spectro-temporal fluctuations critical for speech intelligibility, suggesting that speech acoustic parameters are encoded in this region (Mukamel et al. 2011; Nourski et al. 2009; Pasley et al. 2012).

More recently using ECoG, Mesgarani et al. (Mesgarani et al. 2014) showed that in the STG, high-gamma activity (75-150 Hz) in response to auditory presentation of phonemes is clustered according to phonetic features such as sonority, nasality and stridency, which remarkably are the same distinctive features defined by linguists (Chomsky and Halle 1968). According to linguistic theories, phonemes are described according to sub-phonemic features that distinguish them or can be shared by them. At the neural level, phonemes with common ‘manner-of-articulation’ (i.e., spectro-temporal patterns, such as stridents /sz∫/) evoked more invariant responses than phonemes with common ‘place-of-articulation’ (such as alveolars /tdszn/) – the phonetic gestures of the speaker. This representational structure of phonemes is also supported by scalp EEG recordings (Khalighinejad et al. 2017, Sankaran et al. 2018). However, electrical activity recorded by ECoG or EEG grids reflects average responses of large neuronal populations and is therefore limited in providing insights into activity patterns of single neurons. Indeed, single-unit activity was previously found to differ from local field potentials (LFPs) in various regions and tasks (Donchin et al. 2001; Ossmy et al. 2015).

Consistent with findings from ECoG and EEG studies, previous single-cell studies on phonetic processing revealed a multidimensional neural representation of phonemes by demonstrating that STG neurons are tuned to subsets of phonemes and have a sparse coding scheme (Creutzfeldt et al. 1989; Chan et al. 2013). However, there is no evidence regarding the functional organization of phonemes at the cellular level.

The current study addresses two main questions regarding spiking activity of STG neurons in response to phonemic stimuli. First, we examined whether the functional organization of phonemes as revealed by single-cell activity is dominated by manner- or by the place-of-articulation dimension. Manner features can be mapped to acoustic properties, therefore providing further support to auditory theories of speech perception. Second, we examined whether the functional organization of phonemes as revealed by single-cells activity matches behavioral responses. To this end, we used the same set of stimuli in both a behavioral experiment with healthy subjects and in the experiment with neurosurgical patients implanted with depth electrodes.

## 2 Materials and Methods

### 2.1 Patients and electrophysiological recording

Data was collected from six patients with pharmacologically intractable epilepsy, implanted with intracranial depth electrodes to identify seizure focus for potential surgical treatment (Mukamel and Fried 2012). Subjects were recruited from two centers (UCLA/Tel-Aviv). Electrode location was based solely on clinical criteria. Each electrode terminated in a set of nine 40-µm platinum– iridium microwires (Fried et al. 1999)—eight active recording wires, referenced to the ninth. Signals from these microwires were recorded at 40 kHz using a 64-channel acquisition system. Before surgery each patient underwent placement of a stereotactic headframe, and then a detailed MR image was obtained using a spoiled-gradient sequence, followed by cerebral angiography. Both anatomical and angiography images were transmitted to a workstation in the operating room, and surgical planning was then performed, with selection of appropriate temporal and extra-temporal targets and appropriate trajectories based on clinical criteria. To verify electrode position, CT scans following electrode implantation were co-registered to the preoperative MRI using Vitrea® (Vital Images Inc.). The patients provided written informed consent to participate in the experiments. The study was approved by and conformed to the guidelines of the Medical Institutional Review Board at UCLA and the Tel-Aviv Sourasky Medical Center (Ichilov hospital).

### 2.2 Stimuli and behavioral task

Stimuli were constructed of either consonant-vowel (CV) pairs, or vowels /a e i ou/ presented in isolation. The consonants in the CV syllables were according to the list in Table 1, and the vowel was set to /a/. Patients were presented with 12 repetitions from each CV pair or vowel, 4 from each speaker, in a random order (ISI = 1 second). The patients were instructed to listen carefully to the syllables. All stimuli were recorded in an anechoic chamber with a RØDE NT2-A microphone and a Metric Halo MIO2882 audio interface, at a sampling rate of 44.1kHz. Stimuli were generated by two male and one female Hebrew speakers. The total number of stimuli was 63 (21 phonemes * 3 speakers). Since some patients were native English speakers and some were native Hebrew speakers, we chose phoneme stimuli that are approximately similar across English and Hebrew (verified in a perceptual task with native English speakers; see Phoneme perception experiment). Length and pitch (by semi-tone intervals) were compared across recorded tokens to choose the most highly comparable stimulus-types. This was done by looking at differences in timeline arrangement, using built-in pitch tracker in a commercial software (Logic Pro X). Further cleaning of noise residues in high-resolution mode was done using Waves X-Noise software. Figure S1 shows an example of the waveform of the syllable /∫a/ (top), with the corresponding spectrogram (bottom), articulated by one of the male speakers.

**Table 1.**
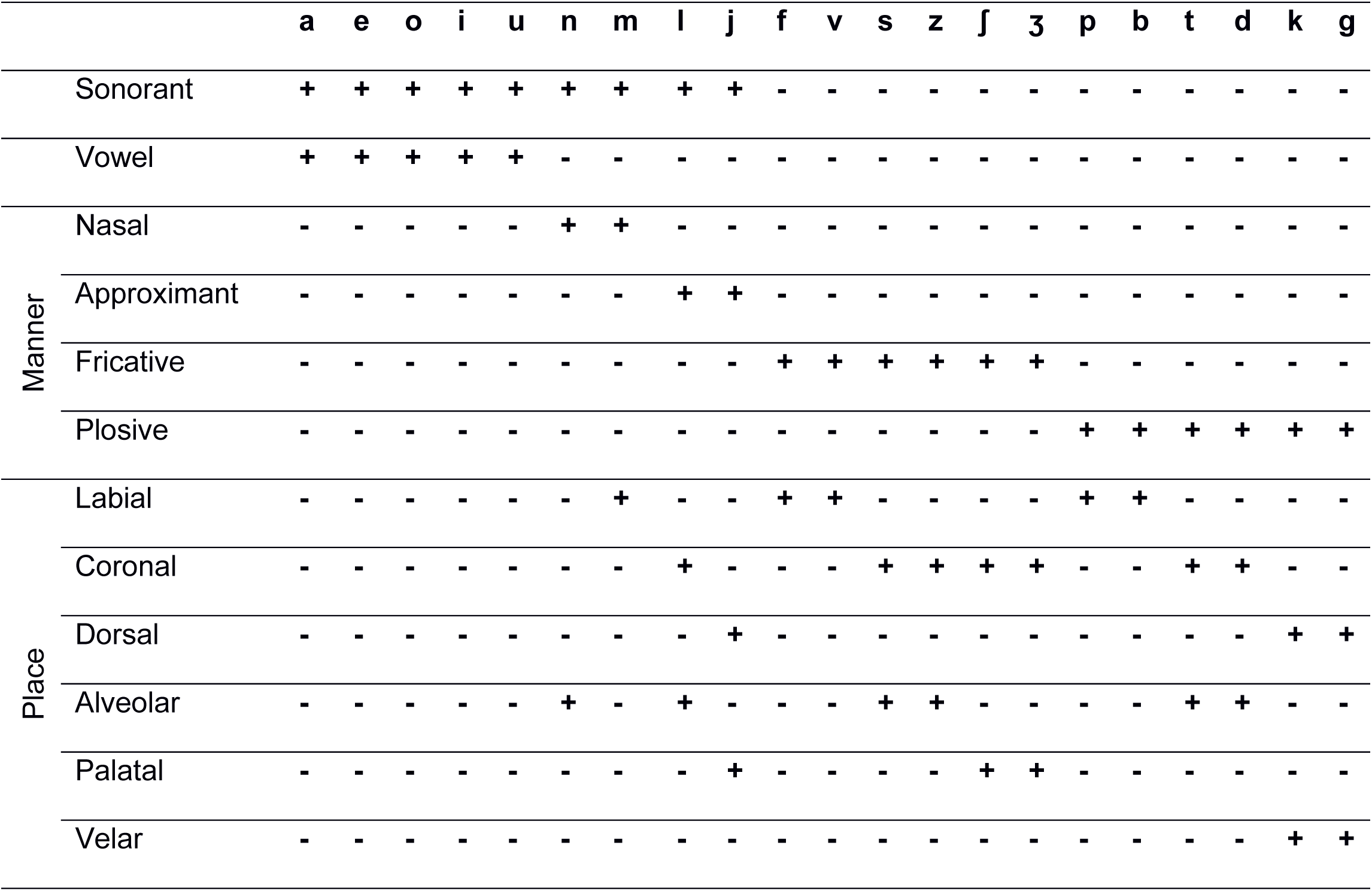
Stimuli details. List of phonemes used in the experiment and their corresponding features.

### 2.3 Data preprocessing

To detect spiking activity, the data was band-pass filtered offline between 300 and 3000 Hz and spike sorting was performed using WaveClus (Quiroga et al. 2004), similar to previous publications (Quiroga et al. 2005; Ossmy et al. 2015). This process allows determining whether the data was recorded from a single- or multi-unit (for full technical details see Quiroga et al. 2005), and yields for each detected unit (single or multi) a vector of time stamps (1ms resolution) during which spikes occurred. To assess responsiveness of each neuron to the phonemes, we computed a t-test between the spike-count distribution before stimulus onset (−500-0ms) and after (0-500ms). Neurons with statistically-significant responses (*p*<0.05) to at least one phoneme were included in subsequent analyses.

### 2.5 Similarity of neural and behavioral responses

To test whether the similarity of phonemes at the behavioral level corresponds with the similarity of population spiking activity in STG, we compared two phoneme similarity matrices - a behavioral and a neural one. The behavioral similarity is calculated from phoneme confusability according to:

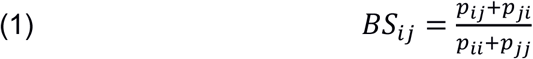

where *p*_*ij*_ is the proportion of times that phoneme i was called phoneme j. *p*_*ii*_ is the hit rate for phoneme i. Thus, *BS*_*ij*_is high, if subjects frequently confused phoneme i with phoneme j (high similarity).

The neural similarity is based on spiking activity in the following way: first, we z-scored the spike-count activity in the response window across all responsive neurons from all patients. Then, for each pair of phonemes i and j, we calculated their Euclidean distance *d*_*ij*_[18] in the response space, and neural similarity was defined according to the following (monotonic) function *NS*_*ij*_ = exp(-*d*_*ij*_).

Finally, we performed Spearman rank correlation between the two matrices. The result is therefore not affected by the exact shape of the function.

### 2.6 Naïve Bayes model

We modeled the observed spike counts from all units assuming that the number of spikes follows a Poisson distribution. Formally, when observed spike-count *x*_*i*_ in unit *i* follows a Poisson distribution *x*_*i*_ *∼ Poisson*(*λ*_*i*_), the probability of observing *k* spikes in a time bin, generated by the unit in response to the presentation of stimulus type *s*, is:

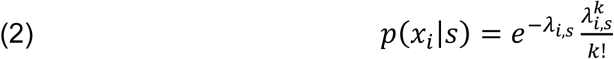

where *λ*_*i,s*_ is the firing rate of unit *i* in response to stimulus type *s*. We modeled the joint spiking activity across units using a Naïve Bayes model. Given a stimulus type (a phoneme or a phonological feature), we assumed that the observed spike counts across units are independent of each other, enabling a simple factorization of the joint probability of stimulus and responses. Formally, the probability of observing a spike-count pattern *x* ∈ N*n* across units in response to the presentation of a stimulus type *s* is:

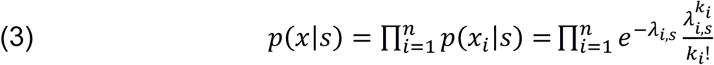

where k_i_ is the number of observed spikes in unit i, and n is the number of units. Random downsampling of the majority classes was performed, in a 5-fold cross-validation procedure. That is, splitting the samples of each class into a training and a test-set according to a 80%-20% ratio, respectively.

### 2.7 Parameter estimation

The parameter estimation of the model is as follow. We estimate the firing rate parameters λ_i,s_ from the training data using maximum likelihood. That is, for each stimulus type s and unit i, we find the firing-rate parameter λ_i,s_ that maximizes the likelihood of observing the spike counts in the training-set trials: 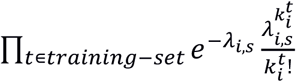, where k_i_ t is the number of observed spikes in unit I in trial t. For the Poisson distribution, as in this case, the maximum-likelihood estimator can be shown to be equal to the mean spike-count.

### 2.8 Inference

Having estimated all firing-rate parameters λ_i,s_, we now describe inference in the model. Given an observed activity pattern across all units x_t_, we infer for each trial t in the test-set the most probable stimulus type s. Using Bayes rule, the posterior distribution is:

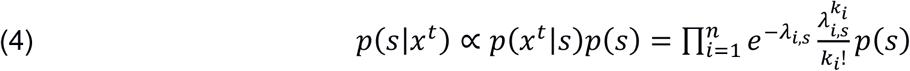

where *p*(*s*) is the prior probability of the stimulus type, which was set as uniform. The mode of the posterior distribution indicates the most probable stimulus type given the firing pattern across units.

### 2.9 Model evaluation

The model is evaluated by comparing the predictions of the model from the inference stage and the ground-truth labels. For binary classification tasks, we use the area under the curve (AUC) as a measure for model performance, with posterior probabilities as scores. For multi-class classification, the full posterior distribution provides additional information compared to its mere mode. For each stimulus type, we calculate the average posterior distribution across all trials in the test set, and use this to construct for each classification task a confusion matrix, in which rows correspond to average posterior distributions. In all cases, statistical significance is determined from the distribution of values across test sets.

## 3 Results

### 3.1 Basic characteristics of the neural responses

We recorded spiking activity from a total of 41 units in six patients implanted with intracranial depth electrodes, while they listened to a variety of phonemes (See Materials and Methods). Of the 41 units, 14 exhibited significant increases in firing rate following stimulus onset and were taken for further analysis (see Materials and Methods and Table 2). Figure 1 depicts rasters and peri-stimulus time histograms (PSTH) plots of spiking activity from one unit in the left STG of one patient. In most neurons, increases in firing rate were observed ∼180ms following stimulus onset, likely due to conductance delays until the signal reaches STG. Some responses contained two activity peaks (e.g., the PSTHs of /b p d s/ in Figure 1) which may be a result of the structure of the stimuli—a consonant followed by the vowel.

**Table 2.** Recording details. Distribution of recorded spiking activity in STG across hemispheres and patients (total responsive STG units = 14).

|  | Left STG | Right STG |
| --- | --- | --- |
| **Patient1** | No units recorded | **Responsive: 3 multi-unit**<br>Not Responsive: None |
| **Patient2** | **Responsive: 1 single-unit; 1 multi-unit**<br>Not Responsive: 1 single-unit; 4 multi-unit | No units recorded |
| **Patient3** | **Responsive: 1 multi-unit**<br>Not Responsive: 1 single-unit; 2 multi-unit | No units recorded |
| **Patient4** | No units recorded | **Responsive: 2 multi-units; 3 single-units**<br>Not Responsive: None |
| **Patient5** | Responsive: None<br>Not Responsive: 2 single unit; 2 multi-unit | **Responsive: 1 multi-unit**<br>Not Responsive: 6 multi-unit |
| **Patient6** | No units recorded | **Responsive: 2 single-unit**<br>Not Responsive: 4 single-unit; 9 multi-unit |

**Figure 1.**
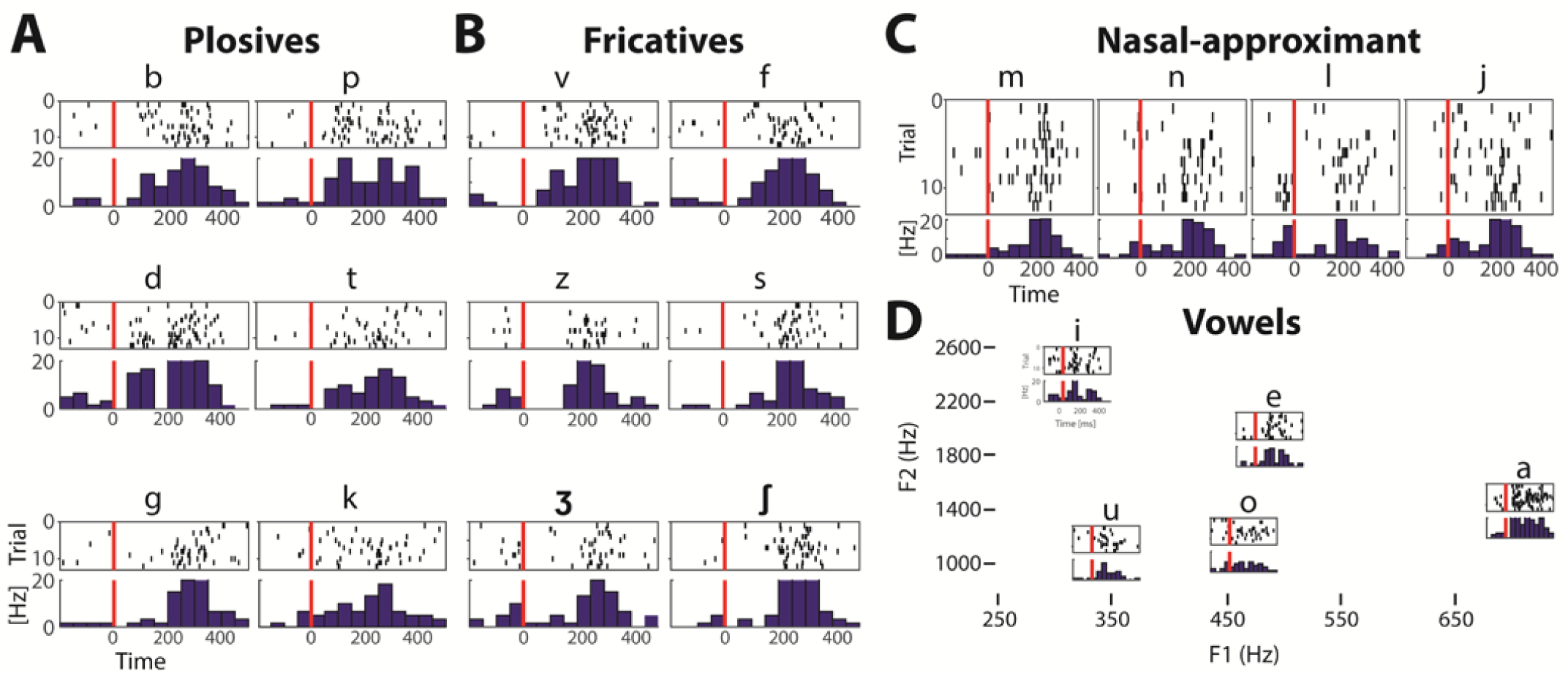
Rasters and Peri-Stimulus Time Histogram plots for one example unit, from patient 3. Consonants are grouped into three groups: plosives, fricatives and nasal-approximant. **(A)** Voiced (left) and unvoiced (right) plosives. **(B)** Voiced (left) and unvoiced (right) fricatives. **(C)** nasal-approximant (left) and affricate (right) phoneme. **(D)** Vowel rasters are embedded in approximate locations in the formant space.

To identify periods for which the neural response is most informative with respect to phoneme identity, we defined a ‘response window’ — the time window for which spiking activity is most separable across phonemes. To that end, we defined a separability index based on the ratio of spike-count variability across trials of different phonemes and trials in which a single phoneme was presented. Spike counts were calculated in 200ms windows, and the separability index was calculated in the range of -100ms to +500ms relative to stimulus onset in steps of 1ms. Figure 2A shows the average of the separability index across all units. The center of the most informative time window is around 180ms after stimulus onset and was used in subsequent analysis (similar to (Chan *et al.* 2013); Changing the time window for calculating spike counts in the range of 100-300ms instead of 200ms did not substantially change the profile of the separability index). This period is similar to the P2 component during phonemic and non-phonemic processing reported in EEG studies, with an activity that peaks at a similar range of time delays from sound onset (Liebenthal *et al.* 2010).

**Figure 2.**
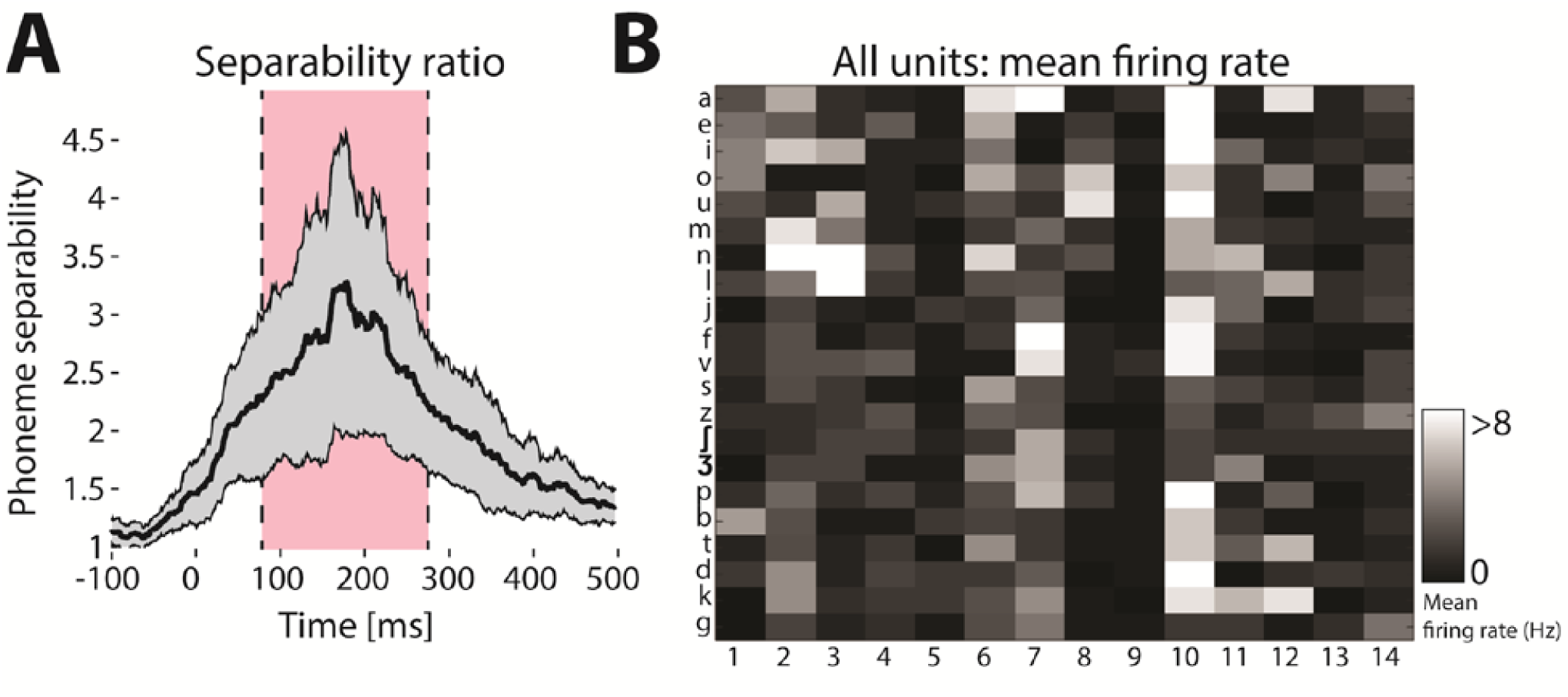
**(A)** Response window. The between-phoneme to within-phoneme variability ratio of the spike-count, for a running window of 200ms (calculated between -100ms to 500ms relative to stimulus onset in steps of 1ms), averaged across responsive units (error bars represent SEM across units). Ticks on the abscissa represent the center of the time window. Response window (79ms-279ms, shaded area) had the maximal value of phoneme separability index. **(B)** Mean firing rates for all units in response to all phoneme stimuli. Color scale represents mean firing-rates for each unit in the response window

### 3.2 The functional organization of phonemes

To examine whether neural responses in STG are functionally clustered, we represented each phoneme as a vector of firing-rate values. To capture the temporal dynamics of the neural response, each phoneme was represented by mean firing rates across trials in four 50ms consecutive bins (Chan *et al.* 2013; Mesgarani *et al.* 2014; Ossmy *et al.* 2015) in the response window (79ms-279ms, see Figure 2) for all fourteen units, giving 56 dimensions in total.

Next, we applied principal component analysis (PCA) to project the neural representation to a lower dimensional space, spanned by two principal components of the data. We found that the sonorant and obstruent phonemes have relatively distinct neural representations, as each group encompasses a different region of the plane (Figure 3A). Based on Euclidean distances among the neural representations of the phonemes, we generated a similarity matrix among the phonemes (Figure 3B, top panel) and performed an unsupervised hierarchical clustering on the similarity matrix. We found a central cluster of obstruents (except for /k/, and including /e/), separated from most sonorants - the vowels /a o i u/ and nasal approximants /n m l j/ (Figure 3B, bottom panel). In addition, the obstruent cluster is further divided into a sub-cluster containing all stridents /s ∫ z 3/. These results point to a functional organization based on manner-of-articulation features, since clustering tends to separate obstruents from sonorants, and to group strident phonemes together. Therefore our next analysis focused on quantifying and comparing response invariances to manner- and place-of-articulation features directly, using a Naïve Bayes model for spike generation (see Materials and Methods for details).

**Figure 3.**
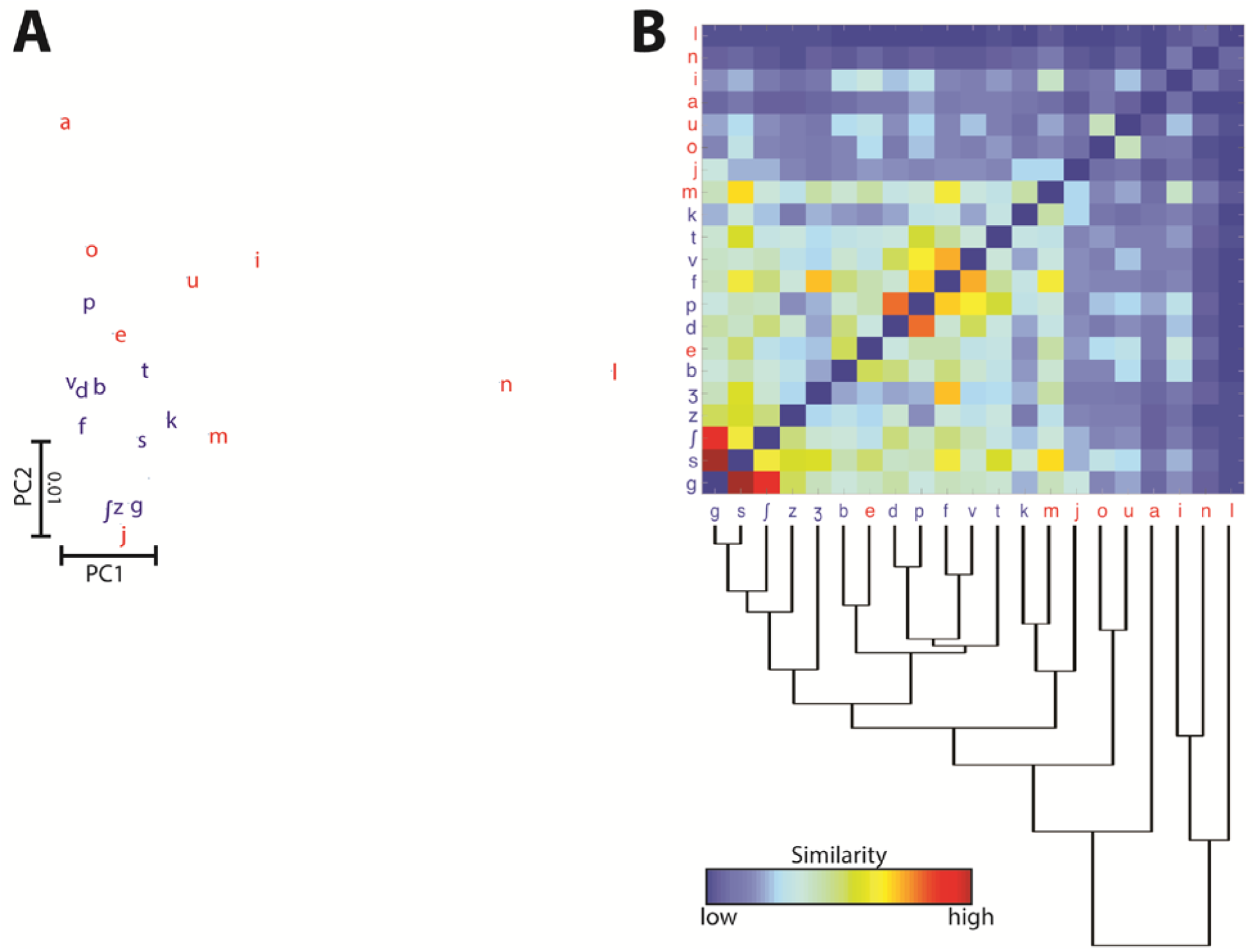
**(A)** Neural representations of phonemes along the first two principal components of the data. Colors: sonorant phonemes (red), obstruent phonemes (blue). **(B)** Hierarchical clustering. Top panel depicts the similarity matrix based on the neural population responses (to enhance color contrasts, diagonal values were manually set to zero). Similarity metric is based on Euclidean distances among the neural representations of the phonemes (see Materials and Methods). Colors: sonorant phonemes (red), obstruent phonemes (blue). Bottom panel depicts hierarchical clustering of the same data.

If manner is a more dominant organizing principle than place, we expect the model to achieve better decoding performance for manner-compared to place-of-articulation features. The confusion errors made by the model are also informative regarding the functional organization of phonemes — higher confusion rate between two classes indicates a higher similarity between their neural representations. If manner is a more dominant dimension at the single-cell level, we expect to observe lower confusion rates of the model among phonemes with different manners of articulation and higher confusion rates among phonemes that share the same manner of articulations.

We examined the performance of the model on two multi-class classifications, for each of the two cases: manner- and place-of-articulation features. For each classification, we labeled the phonemes according to the corresponding phonological features. For manner, we label /a e i o u/ as ‘vowel’, /n m l j/ as ‘nasal-approximant’, /f v s z ∫ 3 / as ‘fricative’, /b d g p k/ as ‘plosives’; and for place-of-articulation, /b p f v m/ as ‘labial’, /t d s z n/ as ‘alveolar’, /∫ 3/ as palatal and /k g/ as velar. We then generated a confusion matrix per classification according to the inferences of the model. Figure 4 shows the significant mean posterior distribution for all phonological features (p < 0.05; t-test compared to chance level), organized in a confusion matrix. Classification according to manner-of-articulation (Figure 4A) resulted in a diagonal structure with higher values on the diagonal, compared to the place-of-articulation classification (Figure 4B). We quantified the extent to which each matrix is diagonal by computing the ratio between the mean of diagonal values and the mean of non-diagonal values. We found a significant difference between the two matrices (manner = 2.89±0.43, place = 1.22±0.49, p < 0.001; t-test).

**Figure 4.**
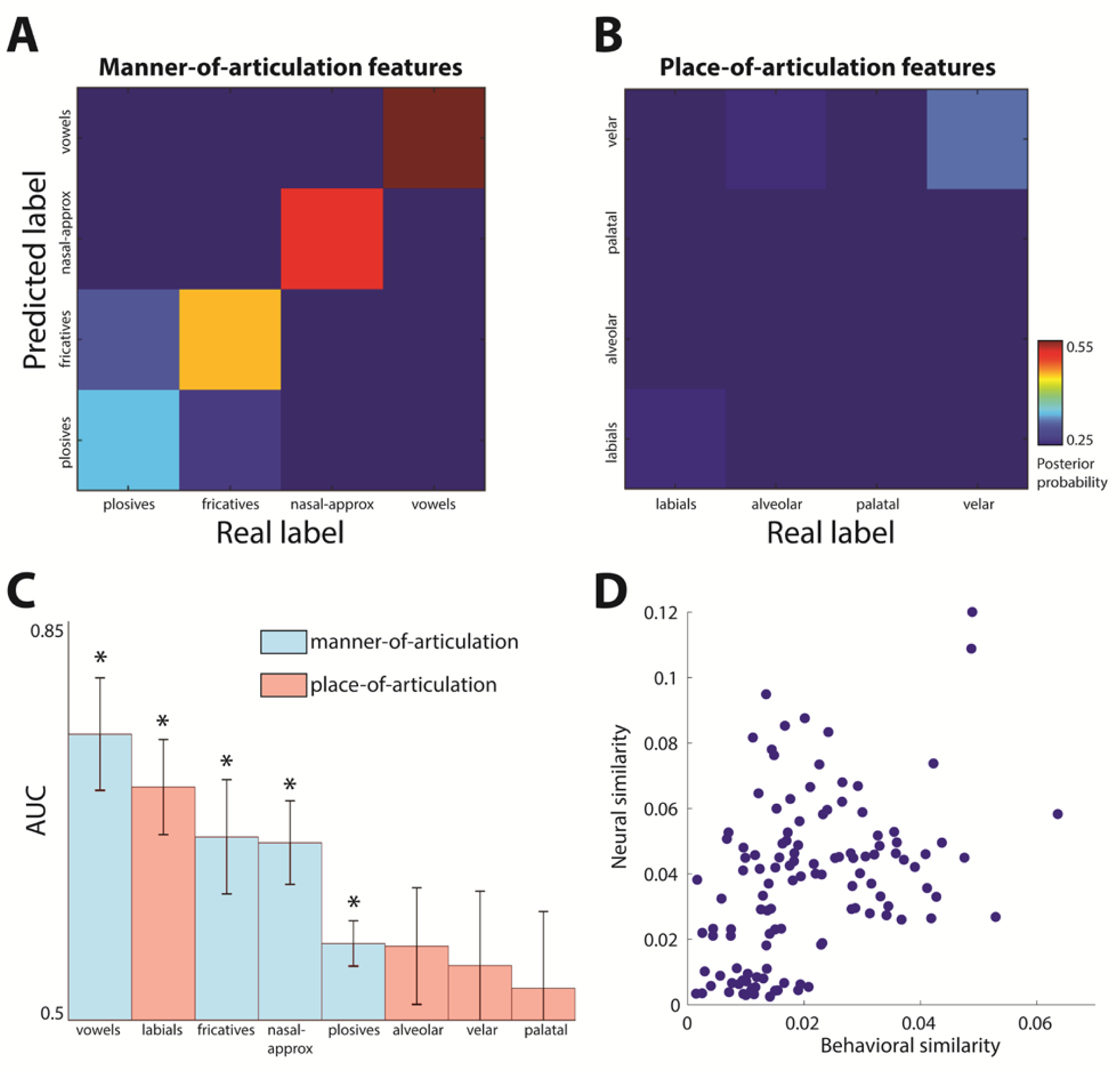
**(A)** Confusion matrix among manner-of-articulation features of consonants: plosives, fricatives, nasals, approximants and vowels. **(B)** Confusion among place-of-articulation features of consonants: labial, alveolar, palatal, velar, glottal (chance level = 0.25). **(C)** AUC values for each binary feature, e.g., [+nasal] vs. [-nasal], [+labial] vs. [-labial]. AUC values were determined from the posterior probabilities of the Naïve-Bayes model and phoneme identities of the test samples; Error-bars are calculated across test sets. **(D)** A comparison between neural and behavioral similarity. Each dot represents a pair of phonemes, X-axis values represent perceptual phoneme similarity, estimated based on confusion rates among phonemes stimuli, which were collected in a behavioral experiment with healthy participants [24]. Yaxis values represent neural similarity from patient data (see Materials and Methods). The Spearman correlation between the behavioral and neural similarities is ρ = 0.45 (*p*<0.001).

To establish the dominance of manner-of-articulation features in distinguishing phonemes, we performed a third classification task. For each phonological feature (e.g., [nasal]), we labeled all phonemes as either + or - ([+nasal] or [-nasal] respectively), and calculated the area under curve (AUC) value for each binary classification. Figure 4C depicts AUC values for all phonological features in descending order. AUC values in all four manner-of-articulation features are significant (p<0.05; compared to chance level, AUC = 0.5) whereas for place-of-articulation, only the labial feature is significantly above chance level.

### 3.3 A comparison between neural and behavioral similarity

Finally, we directly compared neural and perceptual similarities of phonemes. Traditionally, perceptual phoneme similarity is estimated using behavioral tasks, assuming that confusion between two phonemes is correlated with perceptual similarity (Miller and Nicely 1955; Tversky 1977; Shepard 1987). We tested whether phoneme similarity, as estimated in a previous behavioral task (Lakretz et al. 2018), is reflected in neural activity in the STG during listening to the same set of phoneme stimuli.

To that end, we generated one behavioral and one neural similarity matrix. The behavioral similarity matrix is estimated from confusion errors made by thirty-two healthy human subjects, and the neural similarity matrix is derived from the neural representations of the STG responses obtained in the neurosurgical subjects (see Materials and Methods). Since behavioral tasks are limited in generating confusions between consonants and vowels, we focused on the confusion between consonant phonemes only (averaged across subjects). We found a significant correlation between the behavioral and the neural similarity matrices (Figure 4D; ρ = 0.45, p< 10-3, Spearman correlation). This finding suggests that perceptual similarity observed in behavioral tasks can be represented at the level of spiking activity of small population of neurons in STG.

### 3.4 Phoneme perception pretest

Some of the patients in the current study were English speakers and some Hebrew speakers. Therefore, we tested the extent to which the subset of phoneme stimuli used in the experiment is indeed similar across English and Hebrew speakers. To that end, we performed a phoneme perception experiment. Eighteen native English speakers (age range 18.2-35, 12 females; monolingual) sat in front of a screen with headphones and listened to the phoneme stimuli used in the study. After each phoneme, subjects were presented with 21 phonemes on the screen and were asked to select the phonemes they heard. In addition, they were asked to rate their level of confidence in the phoneme selection. Order of played phonemes and options on the screen were randomized across subjects. Subjects identified the phonemes with 79% accuracy (p<0.05; t-test compared to chance level) and high confidence levels (9.2/10 averaged across subjects). Therefore, it is unlikely that differences in native language affected the rest of the results.

## 4 Discussion

A fundamental question in the research of speech perception concerns the functional representation of phonemes in the auditory cortex. Whereas the motor theory of speech perception (Liberman *et al.* 1967; Liberman and Mattingly 1985) suggests that phonemes are organized in terms of common articulatory gestures that generate them (e.g. lip rounding, or jaw raising), auditory theories (Jakobson *et al.* 1951; Stevens 1972, 1989, 2002) argue that phonetic processing is organized based on common spectro-temporal patterns in phoneme waveforms.

Recently, this controversy has been addressed by studies in neuroscience using invasive electrophysiological recordings (Mesgarani *et al.* 2014). These recordings provide a precious glimpse into the neural representations of linguistic entities, such as the objects of speech perception, with high temporal resolution and spatial localization compared to non-invasive recording techniques. Invasive techniques can record extracellular electrical activity either at the level of LFPs or at the level of action potentials generated by single cells (Mukamel and Fried 2012). So far, evidence from invasive recordings regarding the representation of phonemes was based on activity of large populations of neurons, thus leaving open the question regarding the representation of phonemes at the single-unit level. We characterized seemingly distributed, yet possibly clustered, response patterns (14 neurons) to different vowels and consonant-vowel syllables. We directly inquired whether in STG, (a) the organization of phoneme representation at the level of single-cell activity is dominated by manner or by the place-of-articulation; and (b) perceptual representation of phonemes at the behavioral level matches the neural representation at the cellular level.

We found that the structure of the neural representations of phonemes in a relatively small population of neurons demonstrates a separation between sonorant and obstruent phonemes. These findings are in agreement with previous ECoG studies (Mesgarani *et al.* 2014) that examined the organization of phonemes in the STG and found that the dominant distinctive features are manner-of-articulation that contribute most for phoneme classification. The sonorant-obstruent distinction can be described with acoustic properties but not with motor properties, as sonorants have a clear acoustic marker of resonance, with regular patterns in their waveform, whereas sonorant and obstruent involve varied articulations.

We also found that most of the sonorant and obstruent phonemes cluster separately and that strident fricatives form a sub-cluster of the obstruent one. Our findings point to a functional organization based on acoustic cues. First sonorants are highly resonant and have identifiable formant structure compared to obstruents. Second, stridents have a clear acoustic footprint, characterized by high intensity and high-frequency energy. These findings are in agreement with Pasley et al. (Pasley *et al.* 2012) who showed that speech waveforms can be reconstructed from LFPs in the lateral STG, suggesting that encoded information in this region is mainly acoustic.

To further quantify the representations of different phonemes, we trained a probabilistic classifier, which mimics the generation process of spikes, as recorded by the units. We then compared model predictions when grouping phonemes according to various phonological features. Findings show that the confusion matrix of manner features is more diagonal compared to place-of-articulation features. Similarly, area under the curve for binary classification for each feature resulted with significant prediction for all four manner features whereas only one place-of-articulation feature was above chance. Taken together, we show that spiking activity of few cells encodes phonemes according to sub-phonemic features that have acoustic correlates, thus providing additional support to auditory theories of speech perception.

Remarkably, spiking activity from a relatively small number of neurons reflected similarities derived from behavioral results, based on phoneme-confusion experiments using the same set of stimuli. The distinct neural representation of nasal and approximant features with respect to other feature classes, corresponded to their relatively distinct perceptual saliency. These results suggest that the perceptual representation of phonemes can be observed at the level of single neurons. Future research should establish the link between behavioral similarity and neural similarity by investigating different population levels and other regions in the auditory processing hierarchy.

## 5 Conclusions

In sum, our results provide first evidence that the organization of speech perception in single STG neurons is more compatible with auditory theories than motor theories and suggest that activity of single neurons in STG might drive perceptual representation of phonemes during behavior.

## Acknowledgments

The authors thank the patients for participating in the study. We also thank M. Tran and G. Kalendar for administrative help and E. Ho, T. Fields, and E. Behnke for technical assistance; This study was supported by the I-CORE Program of the Planning and Budgeting Committee (grant No. 51/11), The Israel Science Foundation (grants No. 1771/13 and 2043/13; R.M.), and The Human Frontiers Science Program (RGP0057/201, Friedmann). The funders had no role in study design, data collection and analysis, decision to publish, or preparation of the manuscript.

## Notes

### Competing Interest Statement

The authors have declared no competing interest.

### Summary of Updates

Further details regarding the pre-processing of single-cell activity

